# Post-transcriptional regulation of IFI16 promotes inflammatory endothelial pathophenotypes observed in pulmonary arterial hypertension

**DOI:** 10.1101/2024.09.19.613988

**Authors:** Rashmi J. Rao, Jimin Yang, Siyi Jiang, Wadih El-Khoury, Neha Hafeez, Satoshi Okawa, Yi Yin Tai, Ying Tang, Yassmin Al Aaraj, John Sembrat, Stephen Y. Chan

**Affiliations:** Center for Pulmonary Vascular Biology and Medicine, Pittsburgh Heart, Lung, Blood, and Vascular Medicine Institute, Division of Cardiology, Department of Medicine, University of Pittsburgh School of Medicine and University of Pittsburgh Medical Center, Pittsburgh, PA, USA; Department of Molecular Biology, Jeonbuk National University, Jeonju, South Korea; Department of Medicine, Hospital of the University of Pennsylvania, Philadelphia, PA, USA; Department of Computational and Systems Biology, University of Pittsburgh School of Medicine and University of Pittsburgh Medical Center, Pittsburgh, PA, USA; McGowan Institute for Regenerative Medicine, University of Pittsburgh School of Medicine and University of Pittsburgh Medical Center, Pittsburgh, PA, USA; Division of Pulmonary, Allergy, Critical Care, and Sleep Medicine, Department of Medicine, University of Pittsburgh School of Medicine, Pittsburgh, PA, USA

**Keywords:** m6A methylation, pulmonary hypertension, inflammation, endothelial

## Abstract

Pulmonary arterial hypertension (PAH) is a progressive disease driven by endothelial cell inflammation and dysfunction, resulting in the pathological remodeling of the pulmonary vasculature. Innate immune activation has been linked to PAH development; however, the regulation, propagation, and reversibility of the induction of inflammation in PAH is poorly understood. Here, we demonstrate a role for interferon inducible protein 16 (IFI16), an innate immune sensor, as a modulator of endothelial inflammation in pulmonary hypertension, utilizing human pulmonary artery endothelial cells (PAECs). Inflammatory stimulus of PAECs with IL-1β up-regulates IFI16 expression, inducing proinflammatory cytokine up-regulation and cellular apoptosis. IFI16 mRNA stability is regulated by post-transcriptional m6A modification, mediated by Wilms’ tumor 1-associated protein (WTAP), a structural stabilizer of the methyltransferase complex, via regulation of m6A methylation of IFI16. Additionally, m6A levels are increased in the peripheral blood mononuclear cells of PAH patients compared to control, indicating that quantifying this epigenetic change in patients may hold potential as a biomarker for disease identification. In summary, our study demonstrates IFI16 mediates inflammatory endothelial pathophenotypes seen in pulmonary arterial hypertension.

## Introduction

Pulmonary hypertension (PH) is a progressive disease with a poor prognosis, driven by both genetic and epigenetic factors. PH is characterized by an elevation in pulmonary arterial pressure and clinically categorized into 5 major groups by the World Symposium on Pulmonary Hypertension (WSPH) [1]. WSPH Group 1 pulmonary arterial hypertension (PAH) is a severe form of PH, which results in wall thickening of pulmonary vasculature [2]. Pathogenic vascular remodeling and vasoconstriction in PAH are driven in part by endothelial dysfunction, defined by increased inflammation [3]. The importance of inflammation in PAH pathogenesis has been established in recent years, with both PH animal models and PAH patients showing high levels of inflammatory cytokines [4]. Either as an initiating trigger and/or second hit, such inflammatory activation can drive pulmonary vascular dysfunction and remodeling, leading to increased pulmonary arterial pressure and right heart failure [5]. More specifically, our understanding of regulation, propagation, and reversibility of inflammation in PAH remains limited.

Currently available therapeutics, such as vasodilators, are not curative, as they are directed against symptoms rather than the fundamental mechanisms of vascular remodeling involved in disease progression. The approval for clinical use of sotatercept, an anti-remodeling agent targeting transforming growth factor (TGF)-β signaling, suggests such targeted therapeutics hold promise [6]. However, these therapies have not been shown to reverse disease and may not be suitable for all PAH patients. Thus, it is crucial to continue to expand the development of disease-modifying drugs.

Emerging data supports the crucial pathogenic role of endothelial inflammation in PAH. Inflammation is thought to result in endothelial injury, thus further promoting early EC dysfunction and contributing to pathologic changes in underlying cell types, such as smooth muscle cells and fibroblasts [7, 8]. While genetic bone morphogenetic protein receptor 2 (BMPR2) mutations are associated with PAH development, a second stimulus, such as inflammation, is important for pathologic remodeling of the pulmonary vasculature [9, 10]. Increased pro-inflammatory cytokines such as interleukin-1β (IL-1β), interleukin-6 (IL-6), and tumor necrosis factor α (TNFα) have been demonstrated to induce EC dysfunction and arteriolar inflammation in PAH [11]. Notably, several studies have established a role for endothelial inflammatory induction in promoting PAH [12, 13]. Interestingly, interferon gamma inducible protein 16 (IFI16), an important component of the inflammatory signaling cascade has been shown to promote endothelial inflammatory phenotypes [14, 15]. IFI16 has also been shown to inhibit endothelial migration, invasion, tube formation and cell cycle progression [16], pathophenotypes consistent with PAH. With the high expression of IFI16 observed particularly in the vascular endothelium, its cell type specificity may suggest a role for IFI16 in early inflammation of endothelial cells in PAH development [16, 17]. Furthermore, a growing body of literature points to IFI16 as an important mediator of inflammation in Sjogren’s syndrome and systemic sclerosis, autoimmune diseases associated with Group 1 PAH [18, 19]. However, the role of IFI16 as an inflammatory activator in PAH remains unknown.

In this study, utilizing both PAH patient samples and experimental PAH models, we observed a marked increase in IFI16 mRNA levels in pulmonary artery endothelial cells (PAECs). Accordingly, IFI16 knockdown inhibited apoptosis, enhances proliferation, and reduces proinflammatory markers in PAECs. Furthermore, we demonstrate that IFI16 mRNA stability is regulated via inflammation-induced post-transcriptional modification by Wilms’ tumor associated protein (WTAP). WTAP promotes m6A methylation of IFI16 transcript, inducing endothelial dysfunction consistent with that seen in PAH. Taken together, our findings demonstrate that IFI16 pathogenic driver of endothelial inflammation, requiring WTAP-mediated m6A methylation to promote IFI16 mRNA stability. We identified the novel WTAP-IFI16 axis as potential therapeutic target to ameliorate endothelial dysfunction observed in PAH.

## Materials and Methods

### Human Subjects

**Supplemental Tables S1 and S2** describe human PAH patients from which lung and plasma samples were collected. For tissue samples derived from Group 1 PAH patients, diagnosis was made by an expert physician, based on the criteria of having an elevated mPAP > 20 mmHg, pulmonary capillary wedge pressure ≤15 mm Hg, and pulmonary vascular resistance ≥ 3 Wood Units (PVR) by right heart catheterization. For PAH cases, the expert physician adjudicated the diagnosis by ruling out left heart disease, hypoxic lung disease, or chronic thromboembolic disease. For control lung samples, human specimens were collected from unused or discarded samples from the Center for Organ Recovery and Education (CORE) via the University of Pittsburgh Pulmonary, Allergy, Critical Care, and Sleep Medicine Lung Biobank and Repository. Peripheral blood mononuclear cells (PBMCs) were isolated after blood collection, as previously described [20]. All experimental procedures involving the use of human reagents and the study of invasive and non-invasive hemodynamics were approved by the Institutional Review Board (IRB STUDY19050364, STUDY19090084) and the Committee for Oversight of Research and Clinical Training Involving Decedents (CORID) at the University of Pittsburgh. Ethical approval for this study and informed consent conformed to the standards of the Declaration of Helsinki.

### Animal Model

#### Monocrotaline-treated rats

At 12 weeks of age, male Sprague-Dawley rats (Charles River) were injected intraperitoneally (60 mg/kg) with monocrotaline or vehicle. Hemodynamic and histologic measurements of pulmonary vascular disease was performed, as we have previously described.

### Cell Culture

Human pulmonary artery endothelial cells (HPAECs, Lonza CC-2530) were cultured in cell-type specific growth media (Lonza) supplemented with growth factors. In each representative cell culture experiment with PAECs, we used human primary PAECs (non-diseased) derived commercially from different donors; Lot numbers: #0000708987 (64Y, M); #000791935 (58Y, M), #23TL113437 (65Y, F). For a single experiment, cells from a single donor were expanded and plated into separate wells. At least n=3 replicates/group (separate wells) were included in a given experiment, with each well harvested separately for analysis. Experiments were repeated at three independent times using three distinct donor PAECs. If all repetitions showed similar statistically significant changes, the most representative single day experiment in triplicate was presented in the main text, with each data point representing a separate well (see *Statistical Analysis* below). The subsequent repetitions are presented separately in the supplemental text. Cells from passage three through ten were utilized for experiments. For IL-1β experiments, cells were exposed to IL- 1β for 24 hours in complete endothelial cell media.

### Transfection

Human PAECs were transfected at about 70% confluency using OptiMEM media (Thermo Fisher Scientific) and Lipofectamine 2000 (Thermo Fisher Scientific) after siRNA concentration validation. Following a 6-8 hour incubation, transfection media was aspirated and replaced with full serum cell growth media. Cells were analyzed 48 hours after transfection. Silencer siRNAs (Dharmacon, Invitrogen) were used for WTAP, IFI16, BMPR2 knockdowns, and negative control (**Supplemental Table S3**).

### RNA isolation, reverse transcription, and quantitative real-time PCR

To isolate total RNA, cells were collected in QIAzol (Qiagen) and extracted with Rneasy Mini Kit (Qiagen), according to manufacturer’s instructions. cDNA was transcribed using the High-Capacity cDNA reverse transcription kit (Thermo Fisher Scientific), according to manufacturer’s instructions. Quantitative RT-PCR was performed on QuantStudio Flex PCR system (Thermo Fisher Scientific) and normalized to ACTB or GAPDH expression using the ΔΔCT method. TaqMan primers purchased from Thermo Fisher Scientific are listed in **Supplemental Table S4.**

### Immunoblotting

Protein concentration was determined with a Pierce BCA protein assay kit (Thermo Fisher Scientific). 15-20 μg of total protein were loaded into 4-15% gradient SDS-PAGE gels (Biorad) and transferred onto polyvinylidene difluoride (PVDF) membranes (Bio-Rad). Membranes were blocked for one hour at room temperature in 5% non-fat milk in Tris-buffered saline with Tween 20. Membranes were incubated with primary antibody, listed in **Supplemental Table S5**, overnight at 4°C. After primary rinsing, membranes were incubated with appropriate HRP-linked secondary antibody for 1 hour at room temperature. Following secondary rinsing, protein bands were visualized with Pierce ECL reagents (Thermo Fisher Scientific) and Biorad ChemiDoc XRS+. Images were quantified using Image Lab software.

### Plasmid construction, lentivirus production, and transduction

DNA plasmid of IFI16 (Addgene, Plasmid #53741) and WTAP (Addgene, Plasmid #35064) were purchased. HEK293T cells were grown in DMEM containing 10% FBS and transfected with lentiviral plasmids with a packaging plasmid system (LV-MAX Lentiviral Packaging Mix, Thermo Fisher Scientific), according to manufacturer’s instructions. Viral particles were harvested 60 hours after transfection, concentrated, and sterile filtered (0.45 μm). PAECs (70% confluency) were transduced with lentiviral vectors or GFP-containing control vector and polybrene (8ug/ml) in antibiotic-free cell-specific media for 60 hours, repeated twice. Experiments were conducted 120 hours after initiated transduction.

### Apoptosis and Proliferation Assays

Apoptosis was measured using the Caspase-Glo 3/7 Assay (Promega) following the manufacturer’s instructions. Chemiluminescence was measured by spectrophotometer and values were normalized to protein concentration as measure by BCA assay. Proliferation was measured using the BrdU Cell Proliferation Assay Kit (Cell Signaling Technology) following the manufacturer’s instructions after overnight serum starvation for cell cycle synchronization. Absorbance at 450 nm was read with a spectrophotometer to assess colorimetric changes.

### Half-life measurement of IFI16 transcripts

Human PAECs were cultured in standard 6 well culture plates to approximately 80% confluence. Actinomycin D (Sigma, A9415) was added at a concentration of 5 mg/ml. After incubation for indicated time points (0,1,2,4,6 hr), the cells were collected, and RNA samples were extracted for reverse transcription and qPCR. Quantitative RT-PCR was performed on QuantStudio Flex PCR system (Thermo Fisher Scientific). mRNA transcription was inhibited with Actinomycin D and the degradation rate of RNA (Kdecay) was estimated by following equation: ln(C/C0)= -Kdecay(t). C0 is the concentration of mRNA at time 0, t is the transcription inhibition time, and C is the mRNA concentration at the time t. Thus, the Kdecay can be derived by the exponential decay fitting of C/C0 versus time t. The half-time (t1/2), which means C/C0 = 50%/100% = 1/2, can be calculated by the following equation: ln (1/2) = -Kdecay(t1/2). Rearrangement of the above equation leads to the mRNA half-life time value, t1/2 = ln (2/Kdecay).

### MeRIP

MeRIP m6A-PCR was performed by using commercially available Magna MeRIP TM m6A kit (Millipore, 17-10499) according to the manufactural protocol, as previously described [21]. The mRNA was purified from human PAECs using Magnic mRNA isolation kit (New England Biolabs, S1550S). In order to obtain the required quantity of RNA, n=3 technical replicates were pooled into one group, three separate times. Thus, each dot presented in the graph represents each biologic replicate. The 2.5 mg of mRNA was fragmented in 1x fragmentation buffer at 94 degree for 4 minutes and immediately stopped by adding 0.5M EDTA. The fragmented mRNA was purified by RNase MiniElute Cleanup kit (QIAGEN,74204), and the fragment sizes were centered on ∼100nt by agarose get validation. 10% fragmented mRNA was removed as input (10% of Input). 500 mL of MeRIP reaction mixture was prepared by adding 395 mL of fragmented RNA in nuclease free water and 5 mL of RNase inhibitor and 100 mL of 5x IP buffer. The MeRIP reaction mixture was added to prepared Magna protein A/G magnetic beads-m6A antibody tube (1.5 mg of anti-m6A antibody per sample), and incubated for 4 hours at 4°C by rotating head over tail and crosslinked. Each MeRIP reaction tube was briefly centrifuged to remove liquid from cap and sides of the tube and then placed on a separator for 1 minute. The supernatant was carefully aspired without disturbing the magnetic beads. The bead pellets were washed by adding 500 mL of cold 1x IP buffer in the total of 3 times. 100 mL of elution buffer was added to the beads and mixed by gently pipetting several times to completely resuspended beads. The beads containing solution were incubated for 2 hours with by rotating head over tail shaking at 4°C. The MeRIP reactions were centrifuged briefly to remove liquid from cap and sides of the tube and placed on a magnetic separator for 1 minute. The supernatant containing eluted RNA fragments was transferred without aspirating the beads to a new 1.5 mL microcentrifuge tube. The eluted RNA was further purified by RNeasy mini kit to yield final 14 mL of RNA elution. The total RNA elution was applied to qRT-pCR to evaluate target gene enrichment using SYBR Green Master Mix reaction system. The quantification of qRT-pCR results was performed with Ct normalization of the anti-m6A sample, and the negative control of mouse IgG to 10% input sample. The PCR primers were used as follows: IFI16-5’UTR DARCH motif-m6A target: Forward 5’- TCCTGCATTTCTGAAGATCTCAAG-3’, Reverse 5’-AGCCACCTGCATACAACCT-3’.

### Immunofluorescent staining and confocal microscopy

OCT-embedded lung tissue was cryosectioned at thickness of 5-7 microns. Sections were mounted onto gelatin-coated histology slides. Sections were rehydrated in PBS for five minutes, fixed in 4% PFA for thirty minutes, and permeabilized in 0.1% Triton-X 100 for fifteen minutes. After blocking in 5% donkey serum and 2% BSA in PBS for one hour at room temperature, sections were incubated in primary antibodies diluted in 2% BSA overnight at 4°C. Complete summary of primary antibodies included in **Supplemental Table 5**. Sections were incubated in AlexaFluor conjugated secondary antibodies (ThermoFisher) at a 1:2000 dilution in 2% BSA for one hour at room temperature. Slides were then imaged using a fluorescence microscope. Small pulmonary vessel that were not associated with bronchial airways were chosen for analysis. Fluorescence intensity and co-localization was quantified with ImageJ (NIH).

### Statistical analysis

All cell culture experiments presented in the Main Figures represent one of three independent experiments conducted in triplicate, with each data point representing a single well from a particular experiment, as previously described [22], unless otherwise specified. Corresponding repeats of each experiment are displayed in the Supplemental Figures. Unpaired student’s *t*-test was used for comparisons between two groups and one-way analysis of variance (ANOVA) with post-hoc Bonferroni test was used for comparison among more than two groups. A p-value less than 0.05 was considered significant. Data are represented as mean ± SEM.

## Results

### IFI16 is elevated PAH and IFI16 regulates endothelial inflammatory pathophenotypes

Given the undefined role of IFI16 in PAH, we measured IFI16 expression in WSPH Group 1 PAH lungs compared with non-PH lungs. IFI16 gene expression was significantly increased in PAH patient lung samples, suggesting clinical relevance **(Fig. 1A, Table S1)**. Given the role of the IL- 1β as an inflammatory cytokine and known inducer of PAH endothelial pathophenotypes, PAECs were cultured and treated with IL-1β to determine if inflammation up-regulates IFI16. There was a robust increase of IFI16 levels under IL-1β treatment **(Fig. 1B, Fig. S1A)**. Moreover, knockdown of BMPR2, where loss-of-function genetic mutations predisposes to hereditary PAH, resulted in increased IFI16 transcript levels, emphasizing the multifactorial regulation of IFI16 expression by both genetic and environmental factors (**Fig. 1C, Fig. S1B**). Clinically, higher plasma levels of interferon-γ (IFN-γ) have been associated with PAH compared to control patients, and in Scleroderma-PAH compared to Scleroderma only patients [23, 24]. Consistent with the inflammatory regulation of IFI16 and the established correlation of IFN-γ and PAH, induction of IFI16 expression was observed in PAECs treated with IFN-γ **(Fig. 1D, Fig. S1C)**. Additionally, as the inflammatory milieu in PAH is complex and includes but is not limited to IL-1β signaling, we sought to determine if IFI16 is involved in the reprogramming of inflammatory cytokines previously implicated in PAH [25]. We demonstrate under siRNA-mediated inhibition of IFI16, endothelial expression of IL-6, NF-κB, and IL-1β is decreased **(Fig. 1E-G, Fig. S1D-G)**. Beyond the regulation of downstream cytokines expression, we also evaluated the role of IFI16 on endothelial dysfunction. Under the knockdown of IFI16, expression of endothelial proinflammatory markers, VCAM1 and ICAM1, was attenuated (**Fig. 1H, Fig. S1H**). Conversely, IFI16 lentiviral overexpression resulted in up-regulated inflammatory marker levels (**Fig. 1I, Fig. S1I-J**). Additionally, knockdown of IFI16 prevented IL-1β-induced apoptosis (**Fig. 1J, Fig. S1K**), while overexpression increased apoptosis (**Fig. 1K, Fig. S1L**). Together, these findings demonstrate IFI16 as a primary regulator of endothelial inflammation and apoptotic death in PAECs – both known and robust key drivers of PAH.

**Figure 1.**
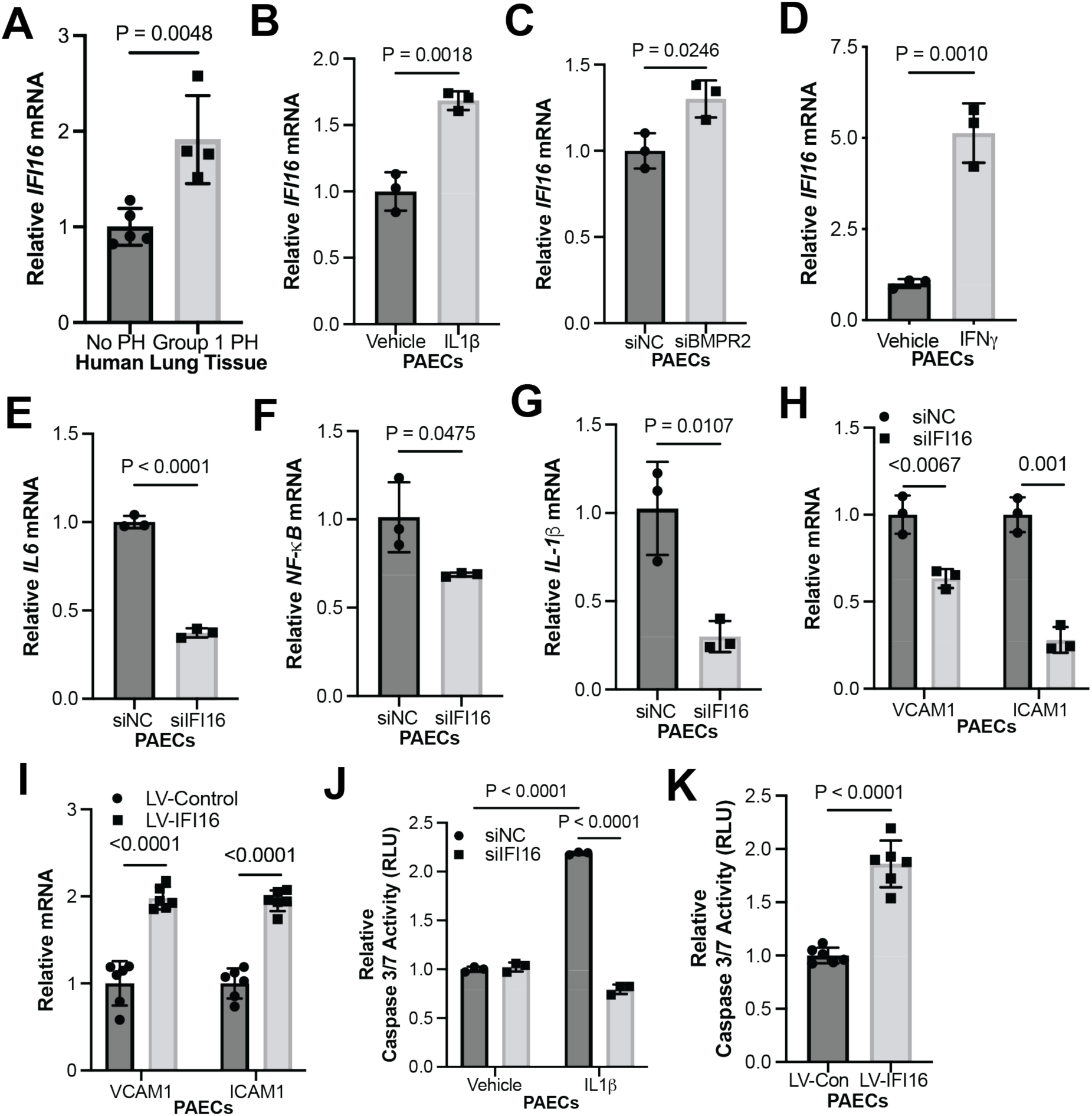
IFI16 is elevated in PAH and promotes endothelial inflammation and apoptosis in human PAECs. (**A**) By RT-PCR, relative IFI16 mRNA was measured in lung tissue from PAH patients (n=4) vs. control (n=5). (**B**) By RT-PCR, relative IFI16 expression (n=3/group) was measured in IL-1β-treated (2 ng/mL) vs. vehicle control (0 ng/mL) PAECs (n=3/group). (**C**) By RT-PCR, relative IFI16 expression was measured following siRNA-mediated BMPR2 knockdown (siBMPR2) vs. control (siNC) PAECs (n=3/group). (**D**) By RT-PCR, relative IFI16 expression was measured in IFNΓ-treated vs. vehicle control PAECs (n=3/group). By RT-PCR, relative expression was of inflammatory cytokines (**E**) IL-6, (**F**) NF-κB, and (**G**) IL-1β were measured in IFI16- deficient (siIFI16) vs. control (siNC) PAECs (n=3/group) (**H-I**) By RT-PCR, relative expression was measured of endothelial inflammatory markers VCAM1 and ICAM1 in IFI16-deficient (siIFI16) vs. control (siNC) PAECs (n=3/group, **H**); conversely, relative expression of these markers was measured following overexpression (LV-IFI16) vs. control (LV-Con) transduction IFI16 in PAECs (n=6/group, **I**). (**J**) Relative caspase 3/7 activity was measured after IFI16 knockdown (siIFI16) vs control in PAECs +/- IL-1β (n=3/group). (**K**) Relative caspase 3/7 activity was measured following IFI16 overexpression (LV-IFI16) vs. control (LV-Con) transduction in PAECs (n=6/group). In all panels, mean expression in control groups was assigned a fold change of 1, to which relevant samples were compared. P-values calculated by two-tailed Student’s *t* test (**A-I, K**), and two-way ANOVA and post-hoc Bonferroni test (**J**), presented as mean +/- SEM.

### Endothelial IFI16 transcript is stabilized by WTAP-mediated m6A methylation under inflammatory conditions

Next, we sought to delineate the regulatory mechanism underlying enhanced IFI16 expression in PAH. Interestingly, recent literature has demonstrated a role for post-transcriptional regulation of innate immune pathway and inflammasome expression via m6A modification of RNA transcripts; this epigenetic mechanism involves the methylation of adenine on RNA transcripts, which subsequently regulates RNA function, including stability, transcription, export and splicing [26, 27]. At the same time, a growing body of literature has defined a role for m6A methylation-dependent smooth muscle cell dysfunction in PH [28–32]. However, the link between IFI16, m6A methylation, and endothelial inflammation has not yet been defined. First, within the IFI16 transcript, we identified a 5’ UTR methylation consensus motif DRACH (D= A, G or T, R= G or A; H= A, C, or U) using the m6A modification site predictor, as previously described [30] (**Fig. 2A**). To evaluate inflammatory regulation of m6A methylation levels, PAECs were treated with IL-1β. Under this inflammatory stimulus, m6A methylation level was enhanced (**Fig. 2B, Fig. S2A**). To identify a candidate m6A regulator mediating increased endothelial m6A methylation, utilizing prior reported microarray profiling of cultured PAECs treated with the cytokine IL-1β vs. vehicle [34], we analyzed the differential expression of known m6A mediators. We found the Wilms tumor associated protein (WTAP) to be the highest up-regulated m6A regulator under IL-1β by transcriptomic analysis (**Fig. 2C**), suggesting a role for WTAP in driving m6A methylation and putative EC pathobiology in PH. In accordance with the observed increase in m6A methylation under pro-inflammatory conditions, in vitro experimentation confirms WTAP RNA up-regulation under IL-1β treatment **(Fig. 2D, Fig. S2B)**, as well as under loss of BMPR2 (**Fig. 2E, Fig. S2C**). Importantly, siRNA-mediated WTAP knockdown reduced total m6A methylation **(Fig. 2F, Fig. S2D-E)**, while lentiviral overexpression of WTAP increased methylation levels **(Fig. 2G, Fig. S2F-G)**, establishing that WTAP controls m6A methylation levels. To establish whether IFI16 expression is mediated by WTAP, we silenced WTAP using two different siRNAs and assessed IFI16 levels, which decreased **(Fig. 2H, Fig. S2H-I)**. Conversely, IFI16 levels increased with WTAP overexpression **(Fig. 2I, Fig. S2J).** To define WTAP regulation of inflammation through IFI16, m6A-specific methylated RNA immunoprecipitation-PCR (MeRIP-PCR) was performed on cultured PAECs, following IL-1β stimulation along with WTAP vs. scrambled control siRNA knockdown. RNA immunoprecipitation was conducted via binding 5’ UTR methylation consensus motif to pull down IFI16 transcript, which was then quantified with RT-PCR. WTAP siRNA-mediated knockdown significantly decreased the IFI16 m6A modification compared to control **(Fig. 2J, Fig. S3)**. Thus, WTAP carries a role in mediating m6A methylation of the 5’ UTR consensus motif of IFI16 RNA transcripts. To investigate how m6A modifications affect IFI16 mRNA stability, PAECs were treated with actinomycin D, an inhibitor of transcription. In that context, WTAP knockdown reduced the rate of exponential decay of IFI16 mRNA dramatically **(Fig. 2K, Fig. S2K)**, demonstrating that m6A methylation of IFI16 transcripts promoted mRNA stability. Overall, these findings establish a mechanism of IFI16 mRNA stabilization via WTAP-dependent post-transcriptional modification in endothelial cells.

**Figure 2.**
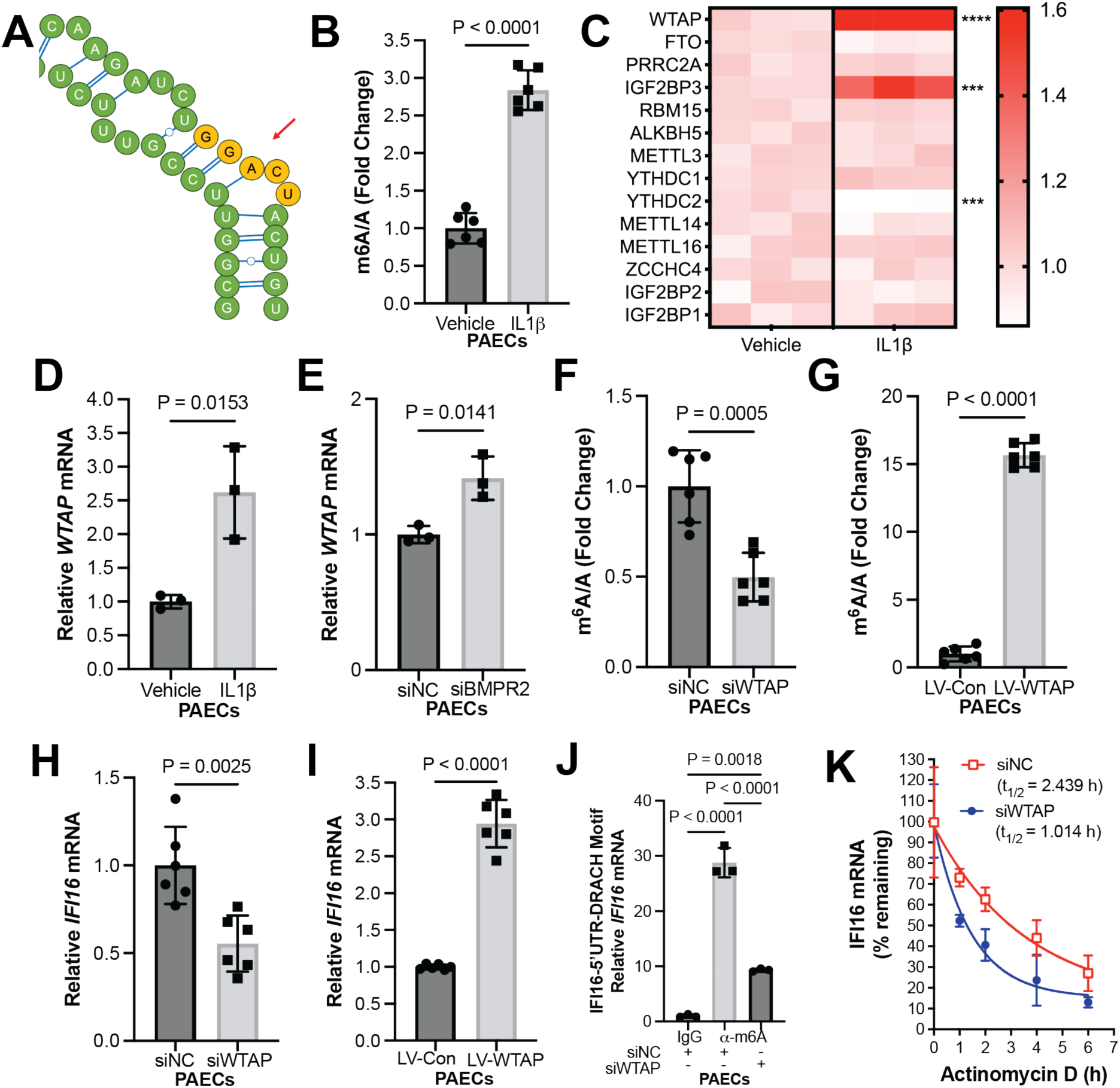
IL-1β promotes WTAP-mediated m6A methylation of IFI16 transcripts, resulting in IFI16 transcript stabilization. (**A**) Schematic of DRACH motif (represented in yellow) of 5’UTR of IFI16 transcript. (**B**) Fold change of m6A/A was measured by colorimetric assay in IL-1β-treated vs. control (vehicle) PAECs (n=6/group). (**C**) A heatmap was generated showing differentially expressed m6A regulators in PAECs treated with or without IL-1β normalized to mean transcript levels in vehicle group, with *** representing p<0.001 and **** representing p<0.0001. (**D**) By RT-PCR, relative WTAP transcript expression was measured (n=3/group) in IL-1β-treated (2 ng/mL) vs. vehicle control (0 ng/mL) PAECs (n=3/group). (**E**) By RT-PCR, relative WTAP expression was measured following siRNA-mediated BMPR2 knockdown (siBMPR2) vs. control (siNC) PAECs (n=3/group). (**F**) Fold change of m6A/A was measured by colorimetric assay in WTAP-deficient (siWTAP) vs. control (siNC) PAECs (n=6/group). (**G**) Fold change of m6A/A was measured by colorimetric assay following WTAP overexpression (LV-WTAP) vs. control (LV-Con) PAECs (n=6/group). (**H**) By RT-PCR, relative IFI16 expression was measured in WTAP-deficient (siWTAP #1) vs. control (siNC) PAECs (n=6/group). (**I**) By RT-PCR, relative IFI16 expression was measured after WTAP overexpression (LV-WTAP) vs. control (LV-Con) transduction in PAECs (n=6/group). (**J**) By MeRIP pulldown (α-m6A vs. IgG control) and RT-PCR, fold enrichment of m6A methylation of IFI16 transcript following IL-1β treatment was measured in WTAP-deficient (siWTAP) vs. control (siNC) PAECs, normalized to IgG negative control. (**K**) IFI16 mRNA decay was measured in WTAP-deficient (siWTAP) vs. control (siNC) PAECs following inhibition of cellular transcription by actinomycin D (n=3/group, presented as mean and 95% CI). In **B-J**, mean expression in control groups was assigned a fold change of 1, to which relevant samples were compared. P-values calculated by two-tailed Student’s *t* test (**B, D-I**) and one-way ANOVA and post-hoc Bonferroni test (**J**), presented as mean +/- SEM.

### WTAP phenocopies IFI16-mediated endothelial dysfunction in human PAECs

Consistent with IFI16-mediated endothelial pathophenotypes, in PAECs, upregulation of proinflammatory markers VCAM1 and ICAM1 was inhibited by WTAP knockdown under IL-1β exposure **(Fig. 3A-B, Fig. S4A-B)** and increased by WTAP overexpression **(Fig. 3C, Fig. S4C)**. Furthermore, WTAP knockdown prevented apoptosis during baseline and IL-1β exposure **(Fig. 3D, Fig. S4D)**, while WTAP overexpression promoted apoptosis in PAECs as measured by caspase 3/7 activity **(Fig. 3E, Fig. S4E)**. Using the monocrotaline (MCT) rat model of PAH, we observe a robust increase in lung WTAP expression, corresponding with increased m6A modifications **(Fig. 3F-H)**. Demonstrating the clinical relevance of these findings, WTAP transcript was increased by RT-PCR in human lung tissue from PAH patients **(Fig. 3I).** Finally, consistent with our lung tissue-specific findings, peripheral blood mononuclear cells (PBMCs) from PAH patients displayed increased levels of m6A methylation **(Fig. 3J)**. Together, these data establish a role for the WTAP-IFI16 axis in inflammatory activation and downstream PAEC apoptosis.

**Figure 3.**
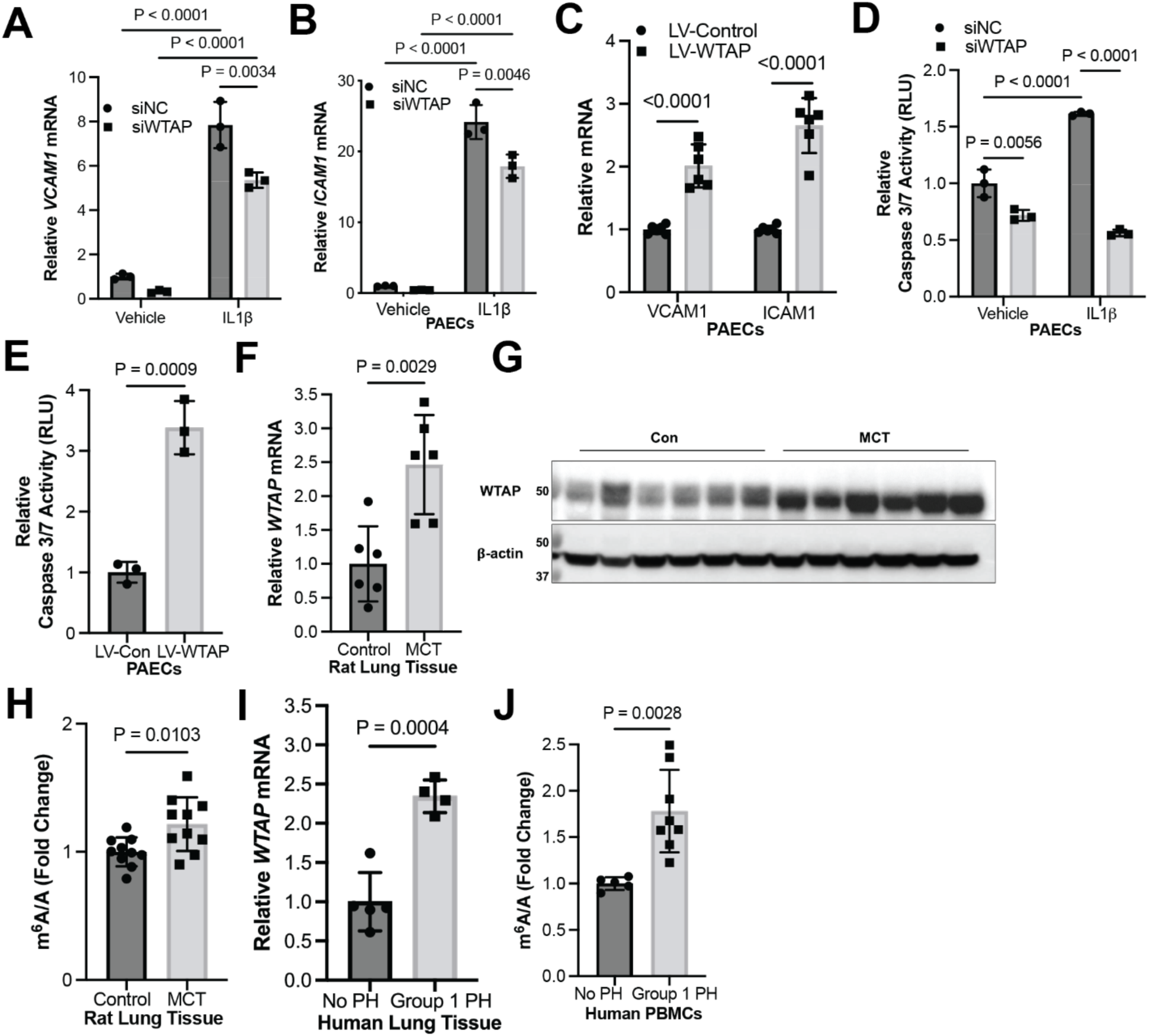
WTAP is elevated in PAH and regulates IFI16-mediated endothelial dysfunction in human PAECs. (**A-B**) By RT-PCR, relative expression was measured of endothelial inflammatory markers (**A**) VCAM1 and (**B**) ICAM1 in WTAP-deficient (siWTAP) vs. control (siNC) with and without IL-1β in PAECs (n=3/group). (**C**) Conversely, relative expression of these markers was measured following WTAP overexpression (LV-WTAP) vs. control (LV-Con) transduction in PAECs (n=6/group). (**D**) Relative caspase 3/7 activity was measured after WTAP knockdown (siWTAP) vs. control in PAECs with and without IL-1β (n=3/group). (**E**) Relative caspase 3/7 activity was measured after WTAP overexpression (LV-WTAP) vs. control (LV-Con) transduction in PAECs (n=3/group). (**F-G**) Relative WTAP transcript expression was measured by RT-PCR (**F**) and WTAP protein expression was measured by immunoblot (**G**) in whole lung from monocrotaline (MCT)-exposed PAH rats (n=6) vs. vehicle control (n=6). (**H**) Fold change of m6A/A was measured in lung tissue from MCT-exposed PAH rats (n=6) vs. vehicle control (n=6). (**I**) By RT-PCR, relative WTAP mRNA was measured in PAH patient lung tissue (n=4) vs. control patients (n=5). (**J**) Fold change of m6A/A was measured in peripheral blood mononuclear cells (PBMCs) from PAH (n=8) vs. control patients (n=5). In all panels, mean expression in control groups was assigned a fold change of 1, to which relevant samples were compared. P-values calculated by two-tailed Student’s *t* test (**C, E, F, H-J**) and two-way ANOVA and post-hoc Bonferroni test (**A, B, D**), presented as mean +/- SEM.

### WTAP and IFI16 are increased in the vascular endothelium in PAH *in vivo*

To determine whether the WTAP-IFI16 is implicated in endothelial cells in an *in vivo* model of PAH, we performed immunofluorescent labeling of IFI16, WTAP, and von-Willebrand factor (vWF), an endothelial cell marker, in MCT-treated rat lungs and vehicle-treated control rat lungs. Consistent with our data in cultured PAECs, IFI16 was up-regulated in MCT rat pulmonary vascular endothelium as compared to control rats (**Fig. 4A-B**). However, total vessel IFI16 signal did not increase significantly (**Fig. 4C**), suggesting that these molecular changes are predominantly implicated in endothelial cells as compared to other pulmonary vascular cell types. Correspondingly, WTAP signal in vWF-positive endothelial cell was enhanced in MCT rat lung compared to control lung (**Fig. 4D-E**), while total vessel WTAP levels were not significantly altered (**Fig. 4F**), supporting endothelial specificity of the proposed mechanism. Thus, the WTAP-IFI16 axis is activated in PAH models both *in vitro* and *in vivo* (**Fig. 5**).

**Figure 4.**
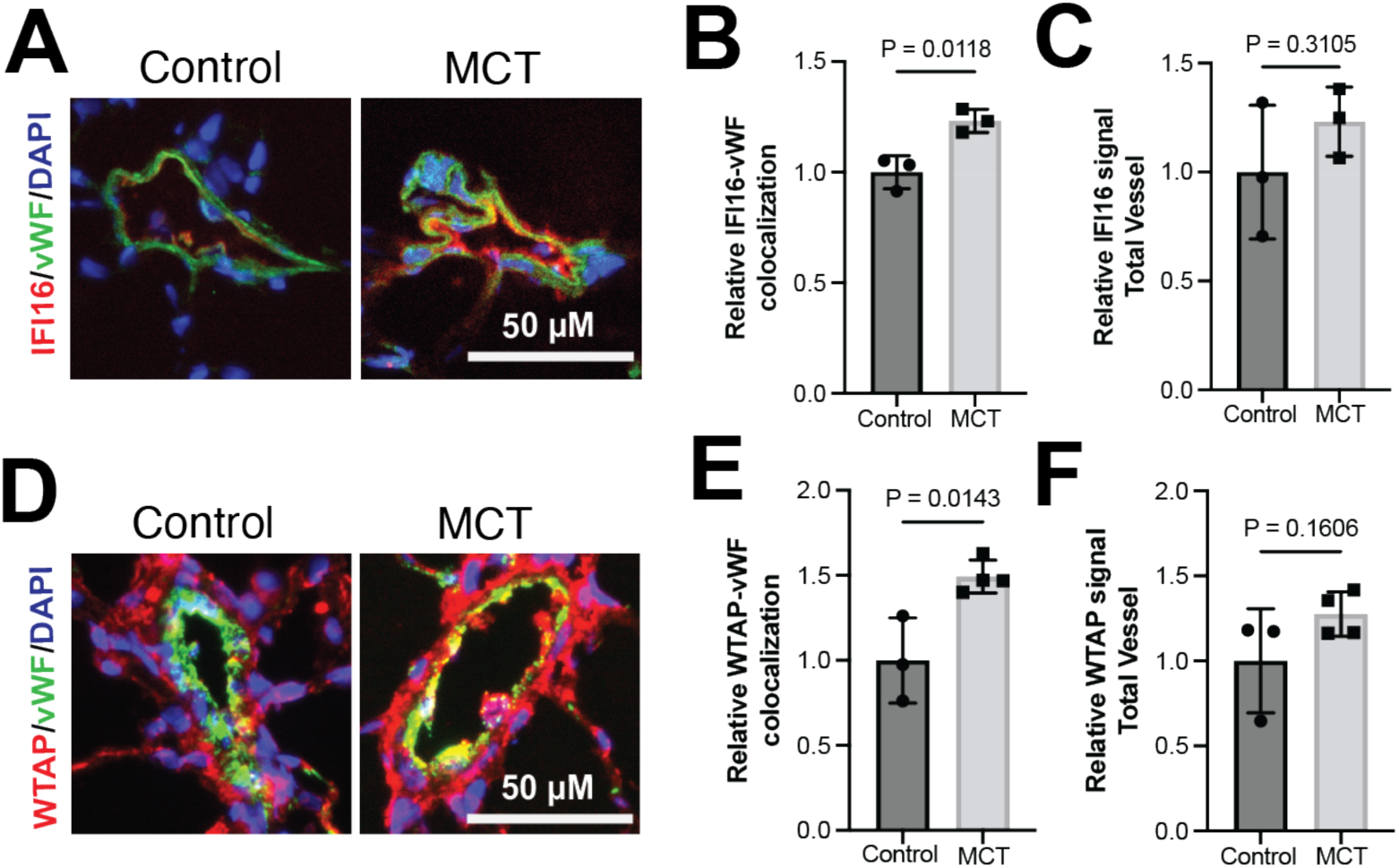
WTAP and IFI16 are upregulated in pulmonary arterial endothelium in PAH *in vivo*. (**A**) Representative images of immunofluorescent staining of IFI16 (red), von Willebrand factor (vWF, endothelial marker, green), and DAPI (blue) in pulmonary vessels, scale bar: 50 µm in vehicle-treated vs. MCT-treated rats. (**B**) Relative IFI16 fluorescence co-localized with vWF-positive signal in vehicle-treated vs. MCT-treated rats. (**C**) Relative IFI16 quantified in total vessel in vehicle-treated vs. MCT-treated rats. (**D**) Representative images of immunofluorescent staining of WTAP (red), von Willebrand factor (vWF, endothelial marker, green), and DAPI (blue) in pulmonary vessels, scale bar: 50 µm in vehicle-treated vs. MCT-treated rats. (**E**) Relative WTAP fluorescence co-localized with vWF-positive signal in vehicle-treated vs. MCT-treated rats. (**F**) Relative WTAP quantified in total vessel in vehicle-treated vs. MCT-treated rats.

**Figure 5.**
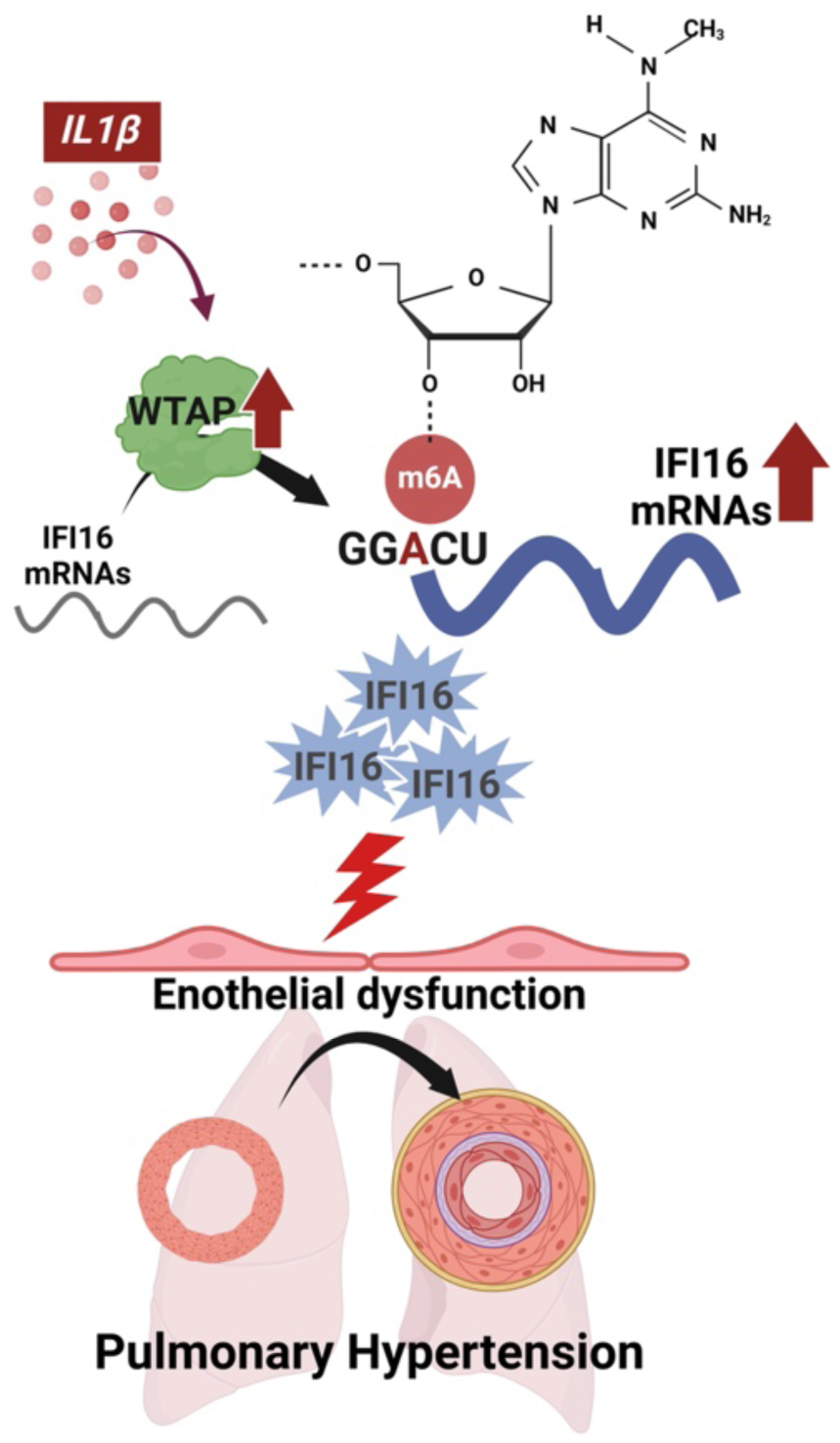
Inflammation promotes m6A methylation-mediated endothelial dysfunction via WTAP-IFI16 axis (model). We demonstrate that IL-1β enhances WTAP activity and downstream methylation of IFI16 mRNA. The resulting up-regulation of IFI16 increases proinflammatory cytokine expression and apoptosis, with implications in PH pathogenesis. IL-1β, interleukin-1β; WTAP, Wilms’ tumor associated protein; and IFI16, interferon gamma inducible protein 16. Images generated by Biorender.com.

## Discussion

Our findings demonstrate that IFI16 is up-regulated in PAECs in human, animal, and cellular models of PAH. Our work supports the model whereby inflammatory-regulated m6A methylation by WTAP controls IFI16 expression and mediates endothelial inflammation and apoptosis (**Fig. 5**). These results identify epigenetic molecular targets that are uniquely attractive for therapeutic development due to the reversible and modifiable nature of m6A methylation. Additionally, our data suggest that alterations of m6A levels in peripheral blood correspond to those seen in PAH lung, thus offering the potential of developing m6A as a blood-borne biomarker of disease.

This study identifies IFI16 as a novel molecular target in endothelial dysfunction. Interestingly, IFI16 is a unique protein that carries two DNA-binding domains and a protein-binding pyrin domain, thus allowing for its versatile function as both a viral DNA sensor and inflammasome inducer, respectively [15]. Moreover, IFI16 is known to localize to both the nucleus and cytoplasm, with additional functions including transcriptional regulation, chromatin remodeling, and extracellular signaling [35]. In the context of endothelial dysfunction, prior studies have found that IFI16 may act as an early driver of the endothelial inflammatory response via its proinflammatory secretome, [17], consistent with our findings indicating IFI16-dependent expression of endothelial proinflammatory markers. As inflammasome induction is a key process required to promote cytokine release to recruit macrophages and T-cells leading to the progression of pulmonary vascular remodeling [36], IFI16 is a promising therapeutic target to prevent endothelial inflammation observed in PAH. However, further studies are necessary to establish the specific mechanism(s) by which IFI16 promotes endothelial dysfunction in the context of PAH, given its diverse functionality.

Additionally, this work furthers our understanding of complex epigenetic regulation of the innate immune system in endothelial inflammation. Highlighting the dynamic regulation of IFI16 expression, a recent study suggests that m6A modification of IFI16 transcripts may facilitate nucleocytoplasmic trafficking, promoting its translation in the setting of viral infection [26], in accordance with our work demonstrating increased RNA stability by m6A modification. Furthermore, WTAP has been implicated in methylating other interferon-regulated transcripts, such as interferon regulatory factor 3 (IRF3). Thus, it has been posited that WTAP may serve as a key mediator of the interferon response, allowing for fine-tuned and modified activation of inflammatory signaling via post-transcriptional m6A regulation [27]. In the context of in PAH pathogenesis, previous literature has emphasized the importance of the hypoxic [28–32], but not inflammatory, regulation of m6A methylation. However, our mechanistic study offers new insights suggesting that not only hypoxic but also inflammatory triggers, such as IL-1β, can activate m6A methylation-mediated pathways via up-regulation of WTAP in *in vitro* PAH models. We demonstrate WTAP further promotes downstream inflammation via stabilization of pro-inflammatory *IFI16*, as evidenced by the up-regulation of endothelial inflammatory markers (VCAM1, ICAM1) and apoptosis in association with increased WTAP, m6A, and IFI16. WTAP and IFI16 deficiency, however, prevented endothelial inflammation. As such, our studies reveal a novel link between m6A post-transcriptional modification and endothelial inflammation observed in PAH, thus setting the stage for future work to determine the complex convergence of hypoxia and inflammation triggers on m6A across pulmonary vascular cell types.

Our work also expands our understanding of the epigenetic landscape in PAH in general, with numerous recent studies reporting roles for DNA methylation, histone modifications, coding and noncoding RNA modifications, and now, m6A mRNA methylation [37, 38]. We hypothesize that these regulatory modifications may be particularly useful in the dynamic regulation of endothelial function, given the barrier role of this cell type in the vasculature and its persistent exposure to toxins, inflammatory cytokines, and hypoxia. Yet, while previous studies have proposed a role for m6A mediators in PAH pathogenesis [28–32], the focus previously has been centered on smooth muscle cell pathobiology. Specifically, m6A modifications have been noted to be the most ubiquitous modification in eukaryotes, with well-established mediators involving three major classes: “writers” (methyltransferases), “erasers” (enzymatic demethylation), and “readers” (regulators of transcript translation, stabilization, and degradation) [39]. Previous work in PAH has suggested a role for writer METTL3/14 in promoting PAH [29], which assembles into an enzymatic complex with structural stabilizer WTAP and other key proteins required for methyltransferase activity [40]. YTH family proteins (readers) have also been established as hypoxic mediators of smooth muscle and endothelial cell function in PAH [30, 32, 41]. Adding to this body of literature, our study demonstrates the role of WTAP, a critical subunit of the RNA methyltransferase complex, in the context of inflammation in PAH models, particularly in endothelial cells and via its regulation of downstream IFI16 RNA transcripts stability. It remains unclear whether WTAP also carries prominent regulatory actions in pulmonary vascular smooth muscle or mural cells and/or whether distinct subsets of m6A regulators may differentially contribute to cell type-specificity in this regard. Taken together, a complex interplay of the writers, readers, and erasers would appear to all regulate PAH, and future studies should be geared toward studying potentially additive or synergistic activities during initiation and progression of PAH in multiple pulmonary vascular cell types.

From a clinical and translational perspective, our work suggests the utility of developing m6A technology in diagnostic, prognostic, and therapeutic applications in PAH. Currently, the capabilities for the early and non-invasive diagnosis of PAH are limited [42, 43]. Given our findings of increased m6A RNA methylation in blood-borne PBMCs in PAH **(Fig. 3J)**, measurement of m6A levels may hold promise for non-invasive detection of PAH. More specifically, with the availability of large PH databases, such as PVRIGoDeep [44] and PVDomics [45], future analyses focusing on comparing m6A modification levels in these cohorts may provide further insight into associations of clinical outcomes with m6A levels. Future studies are also warranted to determine the generalizability of these findings across timing and severity of disease stage and particularly PAH subtype, where autoimmune subtypes of disease may be particularly affected. Furthermore, patients with Sjogren’s syndrome and systemic sclerosis display increased anti-IFI16 autoantibody titers [16]. Thus, our data may be particularly relevant for autoimmune disease and connective tissue disease-associated PAH, warranting further investigation to evaluate IFI16 antibodies as potential biomarkers of disease. Additionally, therapies that reduce m6A methylation could represent an attractive direction within the context of personalized medicine. However, because of the broad extent of m6A methylation across the transcriptome that accounts for nearly 98% of cellular RNA epigenetic activity [43], non-specific therapeutic targeting of m6A regulators such as WTAP carry a substantial risk of “off-target” effects. Alternatively, drugs targeting multiple steps of inflammatory activation are undergoing investigation as clinical PAH therapeutics [12]. In fact, current therapies for PH including prostacyclin analogs and endothelin receptor antagonists, have been shown to target endothelial inflammation [46, 47]. For example, prostacyclin analogs have been shown to decrease VCAM1 levels in systemic circulation [44]. Additionally, bosentan, an endothelin-1 receptor antagonist, was shown to reduce ICAM1 and IL6 levels in systemic circulation and exerts protective effects in PH [48]. Taken in the context of the anti-inflammatory properties of approved PH therapeutics and given the potential for pharmacologic reversibility of m6A methylation of *IFI16*, IFI16 could represent a specific and attractive therapeutic target to reduce endothelial inflammation.

This study has limitations. While the WTAP-IFI16 axis holds biologic activity in controlling endothelial inflammatory phenotypes, *in vivo* studies will be necessary to validate its role in promoting pathologic changes to the pulmonary vasculature. We also did not study the putative roles of other m6A regulators in endothelium and in PAH, where those complexities may dictate a more complex and dynamic profile of m6A regulation on *IFI16* and other transcripts in health and disease. Finally, while our work focused on pulmonary vascular endothelium, we did not include complementary analyses of endothelial cells from various peripheral vascular beds, and thus we do not know the generalizability of these findings across the peripheral vasculature.

In summary, our study defines a mechanistic link between IFI16 induction, m6A methylation by WTAP, and endothelial inflammation consistent with pathophenotypes observed in PAH. Thus, the WTAP-IFI16 axis may represent a compelling translational target in the development of targeted diagnostics and therapeutics in PAH.

## Acknowledgements

This work was supported by American Heart Association Scholarship in Cardiovascular Disease (to R.J.R); R01 HL124021, HL122596, HL151228 (to S.Y.C.). We thank the University of Pittsburgh HSCRF Genomics Research Core (RRID: SCR_018301; Deborah Hollingshead, William Horne, and Janette Lamb) for transcriptomic analyses. We thank the Unified Flow Core at the University of Pittsburgh for use of their instruments. We thank the University of Pittsburgh Pulmonary, Allergy, Critical Care, and Sleep Medicine Lung Biobank and Repository (TTR), the Center for Organ Recovery and Education (CORE), the organ donors, and their families for the tissue samples provided for this work.

## Author Contributions

R.J.R.: Conceptualization, Investigation, Formal Analysis, Visualization, Writing-Original Draft; J.Y.: Conceptualization, Investigation, Formal Analysis, Visualization, Writing-Review & Editing; S.J.: Investigation, Formal Analysis, Visualization, Writing-Review & Editing, W.K.: Investigation, Formal Analysis, Visualization, Writing-Review & Editing, N.H.: Investigation, Formal Analysis, Visualization, Writing-Review & Editing, Y.Y.T: Investigation, Resources, Writing-Review & Editing; Y.T.: Resources, Writing-Review & Editing; Y.A.: Resources, Writing–Review & Editing; S.Y.C.: Conceptualization, Supervision, Project administration, Funding acquisition, Writing- Original Draft.

