## Supplemental Material for "Post-transcriptional regulation of IFI16 promotes inflammatory endothelial pathophenotypes observed in pulmonary arterial hypertension"

**Table S1. Clinical Characteristics for Group 1 PAH Patients (Human Lung Tissue)**

| Group | Age | Gender | mPAP |
|---|---|---|---|
| Control 1 | 36 | M | - |
| Control 2 | 37 | M | - |
| Control 3 | 66 | M | - |
| Control 4 | 13 | M | - |
| Control 5 | 44 | F | - |
| PH 1: Connective tissue disease | 53 | F | 53 |
| PH 2: IPAH | 21 | M | 69 |
| PH 3: IPAH | 50 | F | 56 |
| PH 4: IPAH | 51 | F | 59 |

**Table S2. Clinical Characteristics for Group 1 PAH Patients (Human Blood Samples)**

| Group | Age | Gender | mPAP |
|---|---|---|---|
| Control 1: No PH | 58 | M | 14 |
| Control 2: No PH | 61 | F | 8 |
| Control 3: Healthy control | 40 | F | - |
| Control 4: Healthy control | 66 | F | - |
| Control 5: Healthy control | 53 | F | - |
| PH 1: IPAH | 64 | M | 53 |
| PH 2: IPAH | 66 | F | 46 |
| PH 3: Connective tissue disease | 62 | F | 22 |
| PH 4: IPAH | 40 | F | 42 |
| PH 5: IPAH | 67 | M | 56 |
| PH 6: IPAH | 63 | M | 59 |
| PH 7: IPAH | 72 | M | 24 |
| PH 8: IPAH | 54 | F | 44 |

**Table S3. Silencer RNA Reagent Information (Thermo Fisher Scientific)**

| Reagent | Catalog # |
|---|---|
| Negative control siRNA (#1) | ON-TARGETplus Non-targeting siRNA(D-001810-01-05) |
| IFI16 siRNA | ON-TARGETplus Human WTAP siRNA (LQ-017323-00-0002) |
| WTAP siRNA (siRNA #1) | ON-TARGETplus Human IFI16 siRNA (LQ-020004-00-0002) |
| BMPR2 siRNA | ON-TARGETplus BMPR2 siRNA (LQ-005309-00-0005) |
| Negative control siRNA (#2) | Silencer AM4611 |
| WTAP siRNA (siRNA #2) | Silencer ID: 44219 |

**Table S4. TaqMan Primers Reagent Information (Thermo Fisher Scientific)**

| Reagent | Catalog # |
|---|---|
| *IFI16* | Hs04987070_m1 |
| *WTAP* | Hs00986757_m1 |

| | |
|---|---|
| *VCAM1* | Hs01003372_m1 |
| *ICAM1* | Hs01109748_m1 |
| *IL6* | Hs00174131_m1 |
| *NF-$\kappa$B* | Hs00765730_m1 |
| *IL-1$\beta$* | Hs01555410_m1 |
| *GAPDH* | Hs02786624_g1 |
| *ACTB* | Hs1060665_g1 |
| Mouse *WTAP* | Mm05666501_g1 |
| Mouse *ACTB* | Mm02619580_g1 |

**Table S5. Antibody Reagent Information**

| Company | Use | Reagent | Catalog # |
|---|---|---|---|
| Proteintech | Western blot | WTAP | 60188-1-Ig |
| Santa Cruz | Western blot | ACTB | sc-47778 |
| Santa Cruz | Immunofluorescence | WTAP | sc-374280 |
| ThermoFisher | Immunofluorescence | IFI16 | PA5120684 |
| Abcam | Immunofluorescence | vWF | ab287962 |
| ThermoFisher | Immunofluorescence | vWF | MA5-14029 |

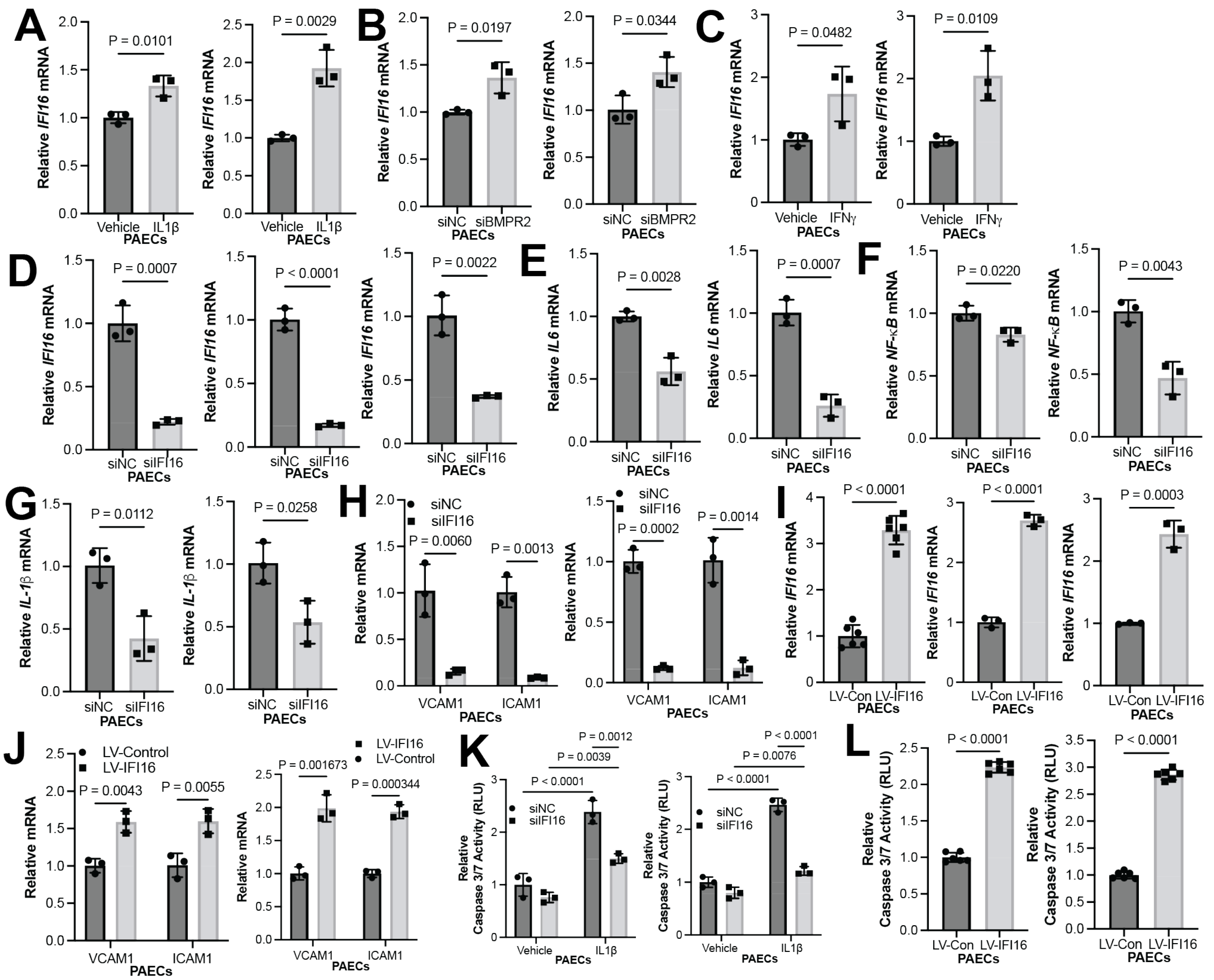


**Figure S1. Inflammatory and genetic regulation of endothelial IFI16 expression.**

(**A**) Experimental repeats of Fig. 1B. By RT-PCR, relative IFI16 expression (n=3/group) was measured in IL-1β-treated (2 ng/mL) vs. vehicle control (0 ng/mL) PAECs (n=3/group). (**B**) Experimental repeats of Fig. 1C. By RT-PCR, relative IFI16 expression was measured following siRNA-mediated BMPR2 knockdown (siBMPR2) vs. control (siNC) PAECs (n=3/group). (**C**) Experimental repeats of Fig. 1D. By RT-PCR, relative IFI16 expression was measured in IFNγ-treated vs. vehicle control PAECs (n=3/group). (**D**) By RT-PCR, relative IFI16 expression was measured in IFI16-deficient (siIFI16) vs. control (siNC) PAECs (n=3/group). Experimental repeats of Fig. 1E-G. By RT-PCR, relative expression was of inflammatory cytokines (**E**) IL-6, (**F**) NF-ΚB, and (**G**) IL-1β were measured in IFI16-deficient (siIFI16) vs. control (siNC) PAECs (n=3/group).

(**H**) Experimental repeats of Fig. 1H. By RT-PCR, relative expression was measured of endothelial inflammatory markers VCAM1 and ICAM1 in IFI16-deficient (siIFI16) vs. control (siNC) PAECs (n=3/group); (**I**) By RT-PCR, relative IFI16 expression was measured following IFI16 overexpression (LV-IFI16) vs. control (LV-Control) PAECs (n=6 or 3/group). (**J**) Experimental repeats of Fig. 1I. Relative expression of VCAM1 and ICAM1 was measured following overexpression (LV-IFI16) vs. control (LV-Con) transduction IFI16 in PAECs (n=3/group). (**K**) Experimental repeats of Fig. 1J. Relative caspase 3/7 activity was measured after IFI16 knockdown (siIFI16) vs. control in PAECs +/- IL-1$\beta$ (n=3/group). (**L**) Experimental repeats of Fig. 1K. Relative caspase 3/7 activity was measured following IFI16 overexpression (LV-IFI16) vs. control (LV-Con) transduction in PAECs (n=6/group). In (**A-L**), mean expression in control groups was assigned a fold change of 1, to which relevant samples were compared. P-values calculated by two-tailed Student's *t* test (**A-J, L**), and two-way ANOVA and post-hoc Bonferroni test (**K**), presented as mean +/- SEM.

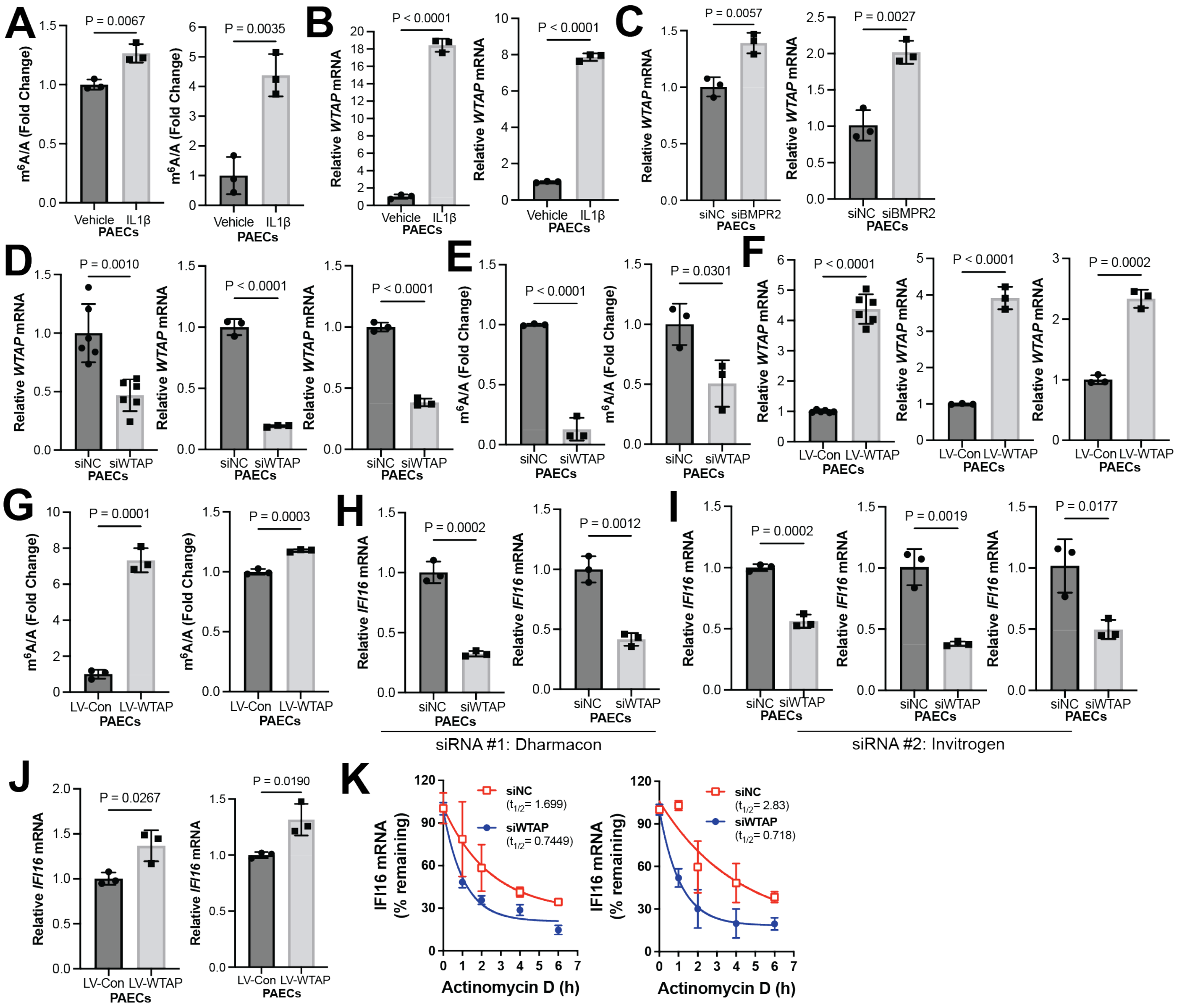


**Figure S2. Inflammatory and genetic regulation of endothelial WTAP expression.** (**A**) Experimental repeats of Fig. 2B. Fold change of m6A/A was measured by colorimetric assay in IL-1β-treated vs. control (vehicle) PAECs (n=3/group). (**B**) Experimental repeats of Fig. 2D. By RT-PCR, relative WTAP transcript expression was measured (n=3/group) in IL-1β-treated (2 ng/mL) vs. vehicle control (0 ng/mL) PAECs (n=3/group). (**C**) Experimental repeats of Fig. 2E. By RT-PCR, relative WTAP expression was measured following siRNA-mediated BMPR2 knockdown (siBMPR2) vs. control (siNC) PAECs (n=3/group). (**D**) By RT-PCR, relative WTAP expression was measured in WTAP-deficient (siWTAP) vs. control (siNC) PAECs (n=6 or

3/group). (**E**) Experimental repeats of Fig. 2F. Fold change of m6A/A was measured by colorimetric assay in WTAP-deficient (siWTAP) vs. control (siNC) PAECs (n=3/group). (**F**) By RT-PCR, relative WTAP expression was measured following WTAP overexpression vs. control PAECs (n=6 or 3/group). (**G**) Experimental repeats of Fig. 2G. Fold change of m6A/A was measured by colorimetric assay following WTAP overexpression (LV-WTAP) vs. control (LV-Con) PAECs (n=3/group). (**H**) Experimental repeats of Fig. 2H. By RT-PCR, relative IFI16 expression was measured in WTAP-deficient (siWTAP) vs. control (siNC) PAECs (n=3/group) using siRNA #1 (Dharmacon). (**I**) By RT-PCR, relative IFI16 expression was measured in WTAP-deficient (siWTAP) vs. control (siNC) PAECs (n=3/group) using siRNA #2 (Invitrogen). (**J**) Experimental repeats of Fig. 2I. By RT-PCR, relative IFI16 expression was measured after WTAP overexpression (LV-WTAP) vs. control (LV-Con) transduction in PAECs (n=3/group). (**K**) Experimental repeats of Fig. 2K. IFI16 mRNA decay was measured in WTAP-deficient (siWTAP) vs. control (siNC) PAECs following inhibition of cellular transcription by actinomycin D (n=3/group, presented as mean and 95% CI). In (**A-J**), mean expression in control groups was assigned a fold change of 1, to which relevant samples were compared. P-values calculated by two-tailed Student's *t* test (**A-J**) presented as mean +/- SEM.

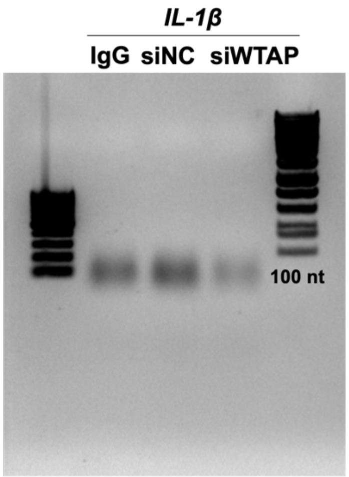


**Figure S3. Fragmentation of RNA for MeRIP.** Calibration of RNA fragmentation and validation of size distribution are shown by gel electrophoresis prior to immunoprecipitation by MeRIP. Fragments converged on ~100 nucleotides in size.

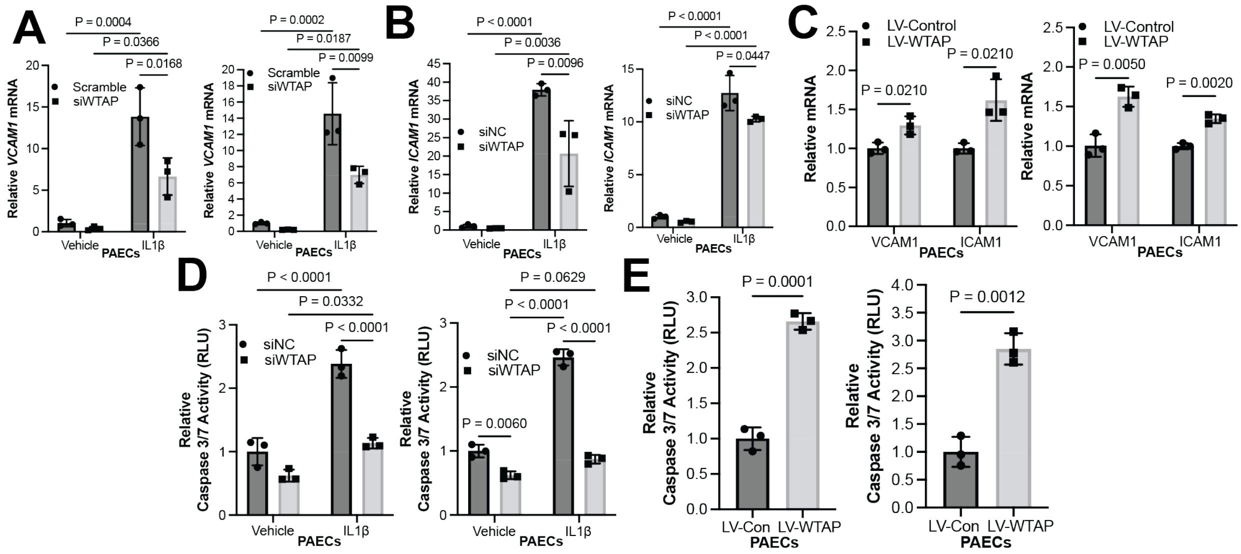


**Figure S4. WTAP phenocopies IFI16-driven endothelial pathophenotypes in PAECs.** (**A-B**) Experimental repeats of Fig. 3A-B. By RT-PCR, relative expression was measured of endothelial inflammatory markers (**A**) VCAM1 and (**B**) ICAM1 in WTAP-deficient (siWTAP) vs. control (siNC) with and without IL-1β in PAECs (n=3/group). (**C**) Experimental repeats of Fig. 3C. Relative expression of VCAM1 and ICAM1 was measured following WTAP overexpression (LV-WTAP) vs. control (LV-Con) transduction in PAECs (n=3/group). (**D**) Experimental repeats of Fig. 3D. Relative caspase 3/7 activity was measured after WTAP knockdown (siWTAP) vs. control in PAECs with and without IL-1β (n=3/group). (**E**) Experimental repeats of Fig. 3E. Relative caspase 3/7 activity was measured after WTAP overexpression (LV-WTAP) vs. control (LV-Con) transduction in PAECs (n=3/group). In all panels, mean expression in control groups was assigned a fold change of 1, to which relevant samples were compared. P-values calculated by two-tailed Student's *t* test (**C, E**) and two-way ANOVA and post-hoc Bonferroni test (**A, B, D**), presented as mean +/- SEM.
